# SVXplorer: Identification of structural variants through overlap of discordant clusters

**DOI:** 10.1101/469981

**Authors:** Kunal Kathuria, Aakrosh Ratan

## Abstract

**Motivation:** The identification of structural variants using short-read data remains challenging. Most approaches ignore signatures of complex variants such as those generated by transposable elements. This can result in lower precision and sensitivity in identification of the more common structural variants such as deletions and duplications.

**Results:** We present SVXplorer, which uses a streamlined sequential approach to integrate discordant paired-end alignments with split-reads and read depth information. We show that it outperforms several existing approaches in both reproducibility and accuracy on real and simulated datasets.

**Availability:** SVXplorer is available at https://github.com/kunalkathuria/SVXplorer.

## Introduction

Structural variants (SVs) that include regions of genomic imbalances called copy number variants (CNVs), and balanced rearrangements such as inversions, account for the majority of varying bases in the human genome. SVs are more common in regions with segmental duplications and have been associated with phenotypes ranging from sensory perception to genomic disorders such as the velocardiofacial and Smith-Margenis syndromes. The discovery and genotyping of these variants remain challenging due to their proximity to repeats, limitations of the alignment algorithms, large non-Gaussian spread in insert size, and the short read lengths typically used in sequencing. SV callers have varying accuracy for different classes of SVs, and some employ specifically designed heuristics for the identification of SV types. However, ignoring signatures of complex SV types often leads to incorrect annotation of common SVs that include deletions, duplications, and inversions. For example, in Fig. 1, ignoring the overlap of signatures from the copy-paste insertion can lead to identification of incorrect breakpoints or the wrong SV types.

**Fig. 1.**
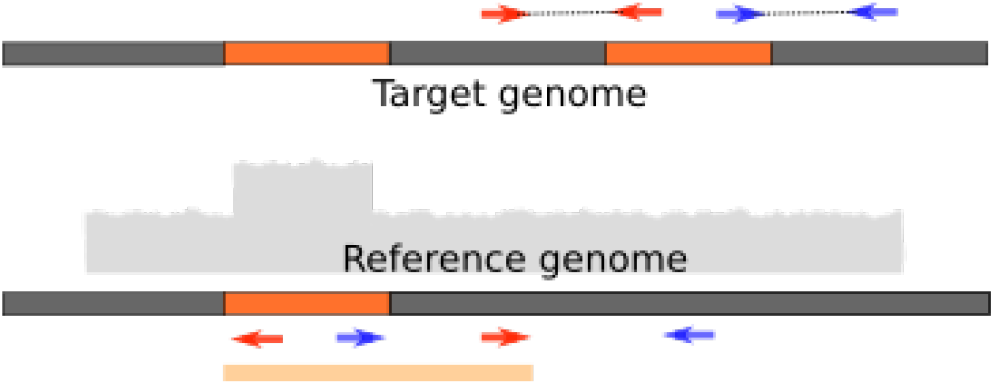
The breakpoints of the duplicated segment might be incorrectly identified as shown at the bottom, if the overlap of signatures from an insertion via ‘copy-paste’ are ignored.

We have developed SVXplorer, which uses a comprehensive 3-tier approach of sequentially using discordant paired-end (PE) alignment, split-read (SR) alignment and read-depth (RD) information to identify multiple SV types while progressively weeding out unlikely candidates. By combining signatures from PE alignment clusters meticulously into “consolidated” variants, integrating and further consolidating PE and SR calls, dynamically calculating PE and SR support thresholds, and corroborating SVs using enhanced local read-depth information, it improves on the precision and sensitivity of calls for the common SV types. Using a combination of probabilistic and combinatorial approaches, SVXplorer shows improvement in comparison to several other popular SV callers on both simulated and real human datasets. On data from two different libraries sequenced from the same cell line, SVXplorer outperforms other methods in both consistency of calls and comparison to calls made using longer PacBio reads. In sequences from a family trio, SVXplorer exhibits the highest fraction of calls that are shared between the child and the parents, while simultaneously identifying the lowest fraction of calls in the child that are not found in either of the parents.

## Methods

SVXplorer requires a coordinate-sorted BAM file generated by aligning Illumina paired-end reads against a reference genome as input. It calculates the coverage and insert length distributions from this BAM file, and groups the fragments that are marked as discordant by the aligner into sets we refer to as clusters. All fragments in a cluster are required to have the same relative orientation of their constituent reads after alignment, and are selected so as to support the same putative variant. It then tests if the clusters can be further grouped into more complex variants such as inversions and translocations based on breakpoint overlap and their combined signature. Split-read evidence from the BAM is then incorporated, both to support existing variants and to create variants that were not captured using the discordant paired-end reads. SVXplorer then processes the variants to remove calls that could be caused due to errors in sequencing or alignment. Finally, read-depth information is added to all the variants and used to further filter the set of calls. We now describe each of these steps in detail.

For clarity, we first define a few terms that are used in the subsequent sections. The “tip” or “head” of an alignment refers to the largest genomic coordinate in case of an alignment to the forward strand, and the smallest genomic coordinate in case of an alignment to the reverse strand of the reference genome. The “tail” analogously refers to the smallest genomic coordinate of a forward-oriented alignment and the largest coordinate for a reverse-stranded alignment. “Mappable” regions refer to regions in the reference that are unlikely to contain reads with poor mapping quality and were identified by running GEM mappability (1) on the reference genome. A “small” cluster refers to a discordant PE cluster that is composed of discordant alignments where the observed insert length is smaller than the estimated mean insert length. A “variant map” refers to the set of all relevant supporting fragments of a putative variant. A “complex” variant is a variant composed of more than one discordant alignment cluster. A “breakpoint region” is the combination of all locations in the reference where the true breakpoint is estimated to possibly exist. A variant whose support tag is “mixed” has support from both PE and SR alignments.

### A. Preprocessing

In this step, we subsample alignments from the input position-sorted BAM file to calculate the insert length and coverage distributions in the dataset. We filter the BAM file to keep discordant reads that pass preset insert length thresholds relative to the mean and respective mapping quality thresholds as input to the next step (see Supplementary Methods for details).

### B. Formation of paired-end clusters

We group fragments aligning discordantly into “clusters” that have the same relative orientation of the reads, and putatively support the same structural variant. Briefly, each fragment with a discordant primary alignment is taken as a node in a graph *G*, and an edge is created between two nodes *i* and *j* if and only if a calculated score *W_ij_* for the pair exceeds a predefined threshold. After all the node pairs in a genomic region have been investigated, connected components from the graph are identified and the nodes in each connected component are separated into maximal cliques using a greedy set-cover approach. Each clique is treated as a set, and the maximum clique (or largest maximal clique) in the collection of cliques, is processed into a cluster, i.e., its member fragments are used to determine the cluster’s breakpoints and error margins. Once a clique is processed, all its member fragments are removed from all other sets, and are not used as part of any other cluster. The clique set itself is now removed from the collection of cliques in the connected component and the steps are repeated. All cliques that have members fewer than a predefined threshold are ignored.

In order to motivate how the score *W_ij_* is calculated, we present a heuristic argument now. Let us define *C_ij_* as the event that two aligned fragments *i* and *j* drawn at random from the genome support the same variant. The connection weight *W_ij_* is a calculated score for the probability of the event *C_ij_*. The distance profile of a pair of fragments *i* and *j*, *D_ij_*, is a function of the difference of the insert length of the two fragments ^1^ and the distance between the respective left reads of the fragments. We denote the observed *difference* in the insert length between the two aligned fragments as Δ*_ij_* and the observed “tip-to-tip” distance between respective left alignments as *L_ij_*. Using Bayes’ rule, *X_ij_* = *P*(*C_ij_|D_ij_* = *d_ij_*) is given by:

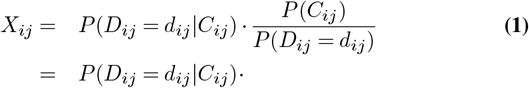

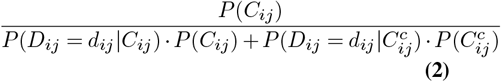

We take note here that the overall probability *P*(*C_ij_*) does not depend on the distance profile, whereas the other terms in Eq. (2) do. We would also like to point out that *P*(*D_ij_* = *d_ij_* | *C_ij_*) is typically a monotonically decreasing function of Δ*_ij_* and *L_ij_*, and 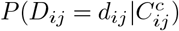 is typically a monotonically increasing function of the same two quantities. The event 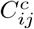 among other things, includes the possibilities that the fragments belong to different variants, or are sampled from systematic misalignments that resemble true variants. Assuming a unimodal insert length distribution and given that alignments clustering together in the reference arising from true variants far outnumber systematic misalignments that cluster together, the above statement should be obvious. In other words, as the difference in insert length between two different fragments with discordant alignments rises, the likelihood of their being sampled from the same genomic region decreases. Further, as the distance between the respective read alignments on either side (e.g., left reads) rises, the likelihood of their belonging to the same variant cluster decreases. It may be more apparent now from Eq. (2) that *X_ij_* is a monotonically decreasing function of Δ*_ij_* and *L_ij_*, as the term multiplying *P*(*C_ij_*) is always less than 1. Also, the only term in Eq. (1) that is grossly dependent on the distance profile is *P*(*D_ij_* = *d_ij_*|*C_ij_*) ^2^. Since the algorithmic objective is to define a fragment-connection weight that is monotonically and structurally similar to *X_ij_*, the following function, a practical reproduction of *P*(*D_ij_* = *d_ij_|C_ij_*), is chosen to define the score between two nodes *i* and *j*:

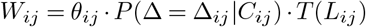

where *P*(Δ = Δ*_ij_|C_ij_*) is directly obtained from the subsampled insert length distribution by binning the insert length difference values, and taking the ratio of the number of entries in the bin in which Δ (the observed insert length difference) resides to the total number of entries. *T*(*L_ij_*) is a function that penalizes distance between the respective left alignment reads after the distance crosses a certain threshold.

The penalty threshold for *T*(*L_ij_*) is chosen to be the “generalized 3 sigma” (*σ*_3_) mark, which is the insert length value at the 99.85 percentile mark (which is equivalent to the 3-sigma mark for Gaussian distributions) of the insert length distribution. The penalty is a simple linear cost that takes *T*(*d*_0_*ij*__) to 0 at *p_mi_*, the insert length at the 99.9999 percentile mark of the insert length distribution. Thus

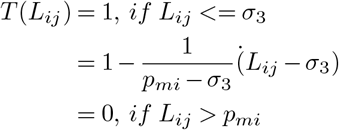

*θ_ij_* is an indicator variable that is 1 if the two fragments (a) have the same relative orientation of reads, and (b) align to the same set of chromosomes. If the relative orientation of the reads is “FR” then they are also required to agree on whether the insert length of the fragments is significantly higher or lower when compared to the average insert length. Currently, a suitable connection weight threshold is applied to the graph: *W_ij_* > 0, i.e., all fragments that have a positive probability of being pairwise connected are connected to each other. However, the overall structure of *W_ij_* is important, as in future work connection weights are envisioned to be edge weights in the graph G, and are to be used in generation of maximal weighted cliques. It is also an important consideration in the regime of low *P*(*C_ij_*), as the structure of *W_ij_* includes hard cutoffs to 0 from discrete sampling of the insert length distribution.

So, in short, fragments are likely candidates for belonging to the same cluster if their mutual insert length difference and their mutual distance are both low, as ascertained from the insert length distribution. The latter is not implied by a mere overlap of the alignment regions if the left and right alignments are distant. After all edges are formed, we find all the maximal cliques of each connected component (2) using an implementation from the Networkx package (3). The cliques are processed into clusters with breakpoints appropriately calculated according to the orientation of the reads. The breakpoint region for each breakpoint of the cluster is given by:

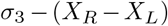

where *X_R_* is the location of the “tip” of the rightmost read supporting the breakpoint, and *X_L_* the location of the “tip” of the leftmost read. This margin offers a conservative estimate even for insert length distribution of anomalous shapes such as those generated when enzyme-based fragmentation methods are used.

### C. Consolidation of paired-end clusters into variants

The clusters that are formed at the end of the previous step are tested for overlap with each other. Cluster “overlap” is defined by overlap of the breakpoint regions in a manner that the composite signature agrees with a specific type of complex variant. Clusters that overlap are grouped and tagged as part of a putative variant. In fact, each cluster is first compared to all such existing variants for possible matches and then to all clusters that are not yet part of complex variants. This allows a variant to be composed of more than two clusters (e.g., translocations). Variant sets are formed by union of the cluster sets described above, recording all the alignments that support a given variant.

Cluster consolidation is detail-intensive, and carefully performed for all basic structural variant (SV) categories that we currently consider. The well-known SV categories used are: deletion, tandem duplication, inversion, de *novo* insertion, and other insertions that occur using a copy- or cut-and-paste mechanisms.

- Deletion (**DEL**): An “FR” cluster that has not been paired with any other cluster and where the included fragments have an insert length that is significantly larger than the average insert length.
- Tandem duplication (**TD**): An “RF” cluster that has not been paired with any other cluster.
- Inversion (**INV**): A pairing of 1 “FF” and 1 “RR” cluster due to the overlap of both left and right alignments respectively.
- Insertion resulting from a copy-paste mechanism (**INS**): A pairing of 1 “FR” and 1 “RF” cluster. An exact signature match as shown in the Fig. S1 is required.
- Insertion resulting from a cut-paste mechanism (**INS_C**): A pairing of 1 “FR” and 1 “RF” cluster as above, but another “FR” deletion cluster flanking 2 adjacent breakpoints (Fig. S2). If all 3 breakpoints lie on the same chromosome (indicating an intrachromosomal translocation), this is a symmetric situation in the 3 breakpoints and it is not possible to distinguish the source of the translocation from the location where it is pasted without using read-depth information. If identified, the paste location breakpoint is defined as “1” and the source locations are defined as “2” and “3”, and the variant is labelled **INS_C_P**.
- De novo insertion (**DN_INS**): A pairing of clusters that are composed of alignments with only one mapped mate and whose alignments have mutually opposite orientation, or an unmatched small “FR” cluster indicating a (novel) inserted segment between its left and right breakpoints.

SVXplorer allows for a detailed treatment of SV types and categories not typically identified using other approaches. Please refer to the Supplementary Methods for a more detailed explanation of these signatures.

### D. Incorporation of split-reads

In this stage, split reads are both used to add support to existing variants and form new variants. Split read alignments (extracted using extractSplitReads_BwaMem script included with LUMPY) are compared to all existing putative variants they could support. If an SR alignment supports a given PE variant call with the correct signature, the variant support tag will now include “SR” and the supporting fragment will be added to the variant map of said variant (see Fig. S5 and Supplementary Methods). If the split alignment does not match any existing (PE or SR) variant, then it is stored as a new possible SR variant. As with PE calls, this new SR variant can be composed of/consolidated by different read signatures, and can be a 2-breakpoint or 3-breakpoint variant.

Variant categories that are created based on SR evidence with no evidence from PE reads are: deletion/insertion, tandem duplication/insertion, insertion and inversion. A brief description of these signatures is provided now, and we include a detailed explanation in the Supplementary Methods.

- Deletion/insertion (**DEL_INS**): A split read yielding unswapped (please refer to Supplementary Methods for detailed explanation of swapping) “FF” or “RR” alignments on the same chromosome is marked as a deletion/insertion candidate. Such a cluster can be supported by both “FF” and “RR” split reads. If this cluster later matches with another cluster, giving rise to a third breakpoint, then it is promoted to an insertion (see Fig. S4). Insertions can be inverted or non-inverted, and depth of coverage is used to disambiguate these calls at a later stage.
- Tandem duplication/insertion (**TD_I**): A split read with the same orientation on the same chromosome that is a swapped read is marked as a tandem duplication/insertion candidate (Fig. S6). Again, it can be promoted to purely an insertion as in the case above. Depth of coverage is later used to disambiguate these cases where possible.
- Insertion (**INS**): Any split read whose segments map to different chromosomes is an insertion candidate. To be counted as a complete insertion, it must match with split reads that create a third breakpoint via the mechanism described above (Fig. S4).
- Inversion (INV): A split read yielding alignments with opposite orientation on the same chromosome is an inversion candidate. To be counted as a complete inversion, an inversion candidate cluster must match with another containing alignments which join the other side of the inversion to the reference.

### E. Variant filtering

For robustness, the statistical procedure used to call variants here is applied to the complete variant set, not to individual clusters.

This procedure was initially conceived for secondary alignments, but works just as effectively if only primary alignments are used. All the variant sets formed thus far can either be completely disjoint or overlap with other variant sets, i.e., share clusters. We require that the *disjointness* of a variant set be the deciding factor in its inclusion in the final variant list. If a variant has a minimum number of supporting alignments (above the mapping quality threshold) that are not shared by any other set then the variant passes this filter.

This disjointness threshold is determined empirically by a simple linear model based on the coverage of the dataset as detailed in the Supplementary Methods. The support threshold set (PE, SR, mixed) for coverage = 25*X*, for example, was (4,4,4).

### F. Incorporation of depth of coverage

This stage carefully assesses all the variant calls using local coverage and filters. Local coverage values in regions between reported variant breakpoints are investigated and if the average coverage in the region seems to contradict the variant in question, then the variant is written as a breakend (BND) event. A BED file listing all mappable regions is recommended as input from the command line and is used to identify regions whose local coverage values can be used in filtering.

In order to calculate variant-region coverage, sampling of bases is done from the middle and edges of the variant region, and only if absolutely necessary, with caveats, from the breakpoint margins. This coverage is assessed relative to the coverage for the chromosome and used to promote potential SR calls to putative variants, or reject PE calls as putative variants. The preset thresholds for deletion and duplication are .8 and 1.2 respectively. If the ratio of average local coverage in the deletion/duplication variant to the chromosomal median coverage (variant coverage ratio, or VCR) exceeds/drops below its respective threshold, then in special cases such variants are not recorded.

Variant calls from all types of clusters (PE, SR, mixed) are rejected if sufficient number of bases (mappable or otherwise) did not exist to calculate coverage in the variant region. Further, coverage is also used break the symmetry of the 3 breakpoints for intrachromosomal translocations and corroborate the source (“cut”) and destination (“paste”) breakpoints.

## Results

We compared SVXplorer (v0.0.3) to several other popular structural variant callers: LUMPY (4), DELLY2 (5), MANTA (6). These algorithms have been used in several large-scale studies including the 1000 Genomes Project, use more than one sources of evidence, and have been shown to be an improvement over most existing tools. We compared their performance on both simulated and real human datasets. LUMPY was run using the defaults in the “lumpy_express” script with the exception of the “-x” option which was used to supply a BED file of regions to be excluded from the analyses. These included regions with abnormally high coverage (4), the mitochondrial genome, the decoy genome and the genome of the Epstein-Barr virus (EBV). DELLY2 was run using the same parameters as used in Layer et al. (mapping quality threshold: 1, minimum support: 4) and an additional BED file with known gaps in the human genome was provided to avoid spurious calls in those regions. MANTA was run using with the default mapping-quality (MQ) threshold and minimum support of 10 and 4 respectively, as in (6). It was provided the same BED file as LUMPY to exclude certain regions that generate unreliable calls. SVXplorer was run with its default parameter set using discordant pairedend (PE) alignments with mapping quality ≥ *1* and split-read (SR) alignments with mapping quality ≥ 10. SVXplorer calculates a minimum support threshold based on the dataset, and was also provided the same exclusion file as LUMPY. In addition a BED file of mappable regions was provided to SVXplorer as input. For all tools, only the variants larger than 100 bps were kept for subsequent analyses.

These specifications were chosen for best overall performance on the human genome for each caller. None of the parameters were changed for any of the callers for any data set, except that no exclude file (or mappable regions file for SVXplorer) was used in processing simulated data. These files are employed only on datasets that involve a human sample (which would likely differ from the reference in such regions and have a high probability of containing misalignments) and not otherwise.

### G. Simulated data

We first ran a haploid simulation wherein RSVSim (7) was used to simulate 2,000 deletions, 1,000 tandem-duplications, 200 inversions, 200 copy-paste insertions and 100 cut-paste insertions (translocations), each of sizes ranging uniformly at random from 100-10,000 bps in the human reference genome (Build 37 decoy), placing breakpoints with a bias towards repeat regions and regions of high homology. We then simulated 100 bp Illumina short-read sequences using wgsim (https://github.com/lh3/wgsim) with a specified mean insert length of 350 and standard deviation of 50, to an average coverage of 50X, and aligned them against the reference genome using BWA mem (8). The four callers were then run on this dataset, and the results were converted to the BEDPE format. The variants were compared to the true breakpoints with a tolerance of 200 bps.

LUMPY, MANTA and DELLY do not identify 3-breakpoint variants such as insertions generated using a cut-paste mechanism, e.g., by DNA transposons, as a single variant, whereas SVXplorer does so. In order to compare tools uniformly, we made relevant adjustments to assess performance. For copy-paste insertions, if a caller identified the two breakpoints of the source location as a “DUP” it counted as a true positive. Cut-paste insertions were identically addressed with “DEL” For SVXplorer, we extracted the source breakpoints from the 3-breakpoint insertion calls and labelled them as either “DUP” or “DEL” according to the insertion type.

Sensitivity and precision was computed for each variant category. The same simulation was repeated at coverages ranging from 2X to 48X in steps of 2X to assess how well the callers perform with varying sequenced information. The relative performance for all callers based on sensitivity of the calls is shown in Fig. 2. As expected, none of the tools made substantial false-positive calls at coverages higher than 6X (Fig. S8), with SVXplorer leading by a small margin over others. SVXplorer has the highest sensitivity for deletions and duplications at all depths of coverage that were investigated. MANTA has the highest sensitivity for inversions closely followed by SVXplorer and DELLY. The default specifications for SVXplorer are conservatively aimed at real data and they mandate that an inversion not be called unless evidence is seen at both ends of the variant. In fact, as will be seen below, the number of inversion calls made by SVxplorer and LUMPY relative to other variants for real datasets are fewer, and much more in line with what is expected.

**Fig. 2.**
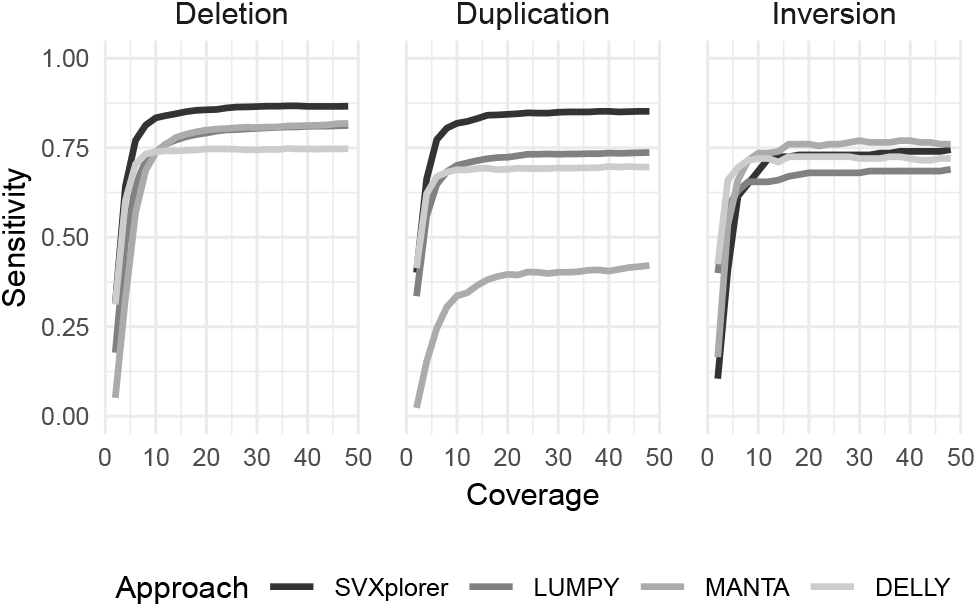
SVXplorer is more sensitive compared to the other approaches even at relatively low genome coverage, as assessed using this simulated dataset.

SVXplorer identifies deletions (duplications) stemming from cut-paste (copy-paste) insertion mechanisms by annotating them as the source breakpoints of the 3-breakpoint insertion calls that pass read-depth filters. The other methods identify them as “FR” (“RF”) clusters that pass coverage filters. As we show in Fig. 1, this can lead to identification of incorrect breakpoints, even if read-depth signature is included in the analysis^3^.

### H. Real data

We next applied SVXplorer along with the other callers to several real human sequencing datasets to evaluate its relative effectiveness under different conditions. Build 37 of the human genome (GRCh37+decoy) was used as the reference for all datasets. For predictive power, the callers were either evaluated against calls made using PacBio long reads, or those made using ensemble approaches such as Parliament (9). Sensitivity, precision and F1 score were computed for all callers after removing calls less than 100 bps from both the call set and the truth set. A call in the “truth” set that overlaps a predicted call within a slop of 200 bps is defined as a true positive. Wherever possible, an assessment was made as to the self-consistency of calls made by each caller for related samples (different libraries or related individuals).

#### H.1. CHM1

Performance was first evaluated on the CHM1 cell line, derived from a human haploid hydatidiform mole. Chaisson et al. (10) released a comprehensive set of structural variant calls that are publicly available at http://eichlerlab.gs.washington.edu/publications/chm1-structural-variation.

As we show in Table 1, SVXplorer outperforms the other methods in sensitivity, precision and F1 score in the identification of deletions.

**Table 1.** Comparison of the various approaches based on CHM1 deletions. The best result for each metric is highlighted in bold.

|  | SVXplorer | LUMPY | DELLY | MANTA |
| --- | --- | --- | --- | --- |
| Calls | 1759 | 1804 | 1116 | 381 |
| True-positives | **1219** | 1174 | 397 | 107 |
| Sensitivity | **27.3** | 26.3 | 8.9 | 2.4 |
| Precision | **69.9** | 65.0 | 35.5 | 28.0 |
| F1-score | **39.3** | 37.5 | 14.2 | 4.4 |

#### H.2. NA12878

We next applied SVXplorer to two separate libraries for the well studied NA12878 cell line (accessions: ERR194147 and SRR505885) along with the other callers.

The callers were evaluated for deletion calls against calls made using PacBio long reads that passed quality filters. For both libraries SVXplorer exhibits better performance characteristics compared to other callers. The results are shown in Table 2. SVXplorer took 51 minutes to process 52X data (ERR194147 accession), which is a fourth of the time taken by the workflow of the other methods. This includes its handling of complex variants and incorporation of split reads meticulously into these complex variants using a hash-based approach.

**Table 2.** Comparison of deletion calls made by the various methods for NA12878. The best result for each metric is highlighted in bold.

| Library |  | SVXplorer | LUMPY | DELLY | MANTA |
| --- | --- | --- | --- | --- | --- |
| ERR194147 | Calls | 2724 | 2607 | 2407 | 787 |
|  | True-positives | **2401** | 2274 | 1867 | 602 |
|  | Sensitivity | **64.9** | 61.5 | 50.5 | 16.3 |
|  | Precision | **88.6** | 87.2 | 77.5 | 76.5 |
|  | F1-score | **74.9** | 72.1 | 61.2 | 26.8 |
| SRR505885 | Calls | 3141 | 3439 | 2407 | 787 |
|  | True-positives | 2541 | **2624** | 1867 | 602 |
|  | Sensitivity | 68.7 | **71.0** | 50.5 | 16.3 |
|  | Precision | **80.7** | 76.2 | 77.5 | 76.5 |
|  | F1-score | **74.2** | 73.4 | 61.2 | 26.8 |

In addition, performance curves for sensitivity and precision with varying coverage were generated for all callers for the ERR194147 library against the PacBio deletion truth set. SVXplorer shows the highest sensitivity and precision even at lower coverage compared to the other callers (Fig. 3). Next, we tested the various callers reproducibility via calls made by each for the two sequencing libraries. We asked the question: “What percentage of calls made by each caller for one library were *found* in the other library”? For this, we take the final call-set of one library (called the “base library”) and compute its overlap with the ungenotyped call set for the other library within a slop of 200 bps. This is because we are interested in knowing whether a variant called in one sample presents reasonable evidence of being seen in the other sample, given various discrepancies and artifacts in fragmentation, sequencing, and alignment. LUMPY, MANTA and DELLY all identify 2-breakpoint variants only, i.e., an “FR” cluster becomes a deletion candidate and an “RF” cluster becomes a duplication candidate. This is true for simple deletions and tandem duplications, but not when clusters with these signatures arrive from cut- or copy-paste insertions. To have the same framework as the other callers, SVXplorer’s self-consistency comparison is done at the cluster level (prior to complex variant formation). Essentially, all its PE and SR clusters that pass filters in the base library are compared to all PE and SR clusters in the other library (with “FR” clusters or equivalents termed “deletions” and “RF” clusters or equivalents termed “duplications” for uniformity across tools).

**Fig. 3.**
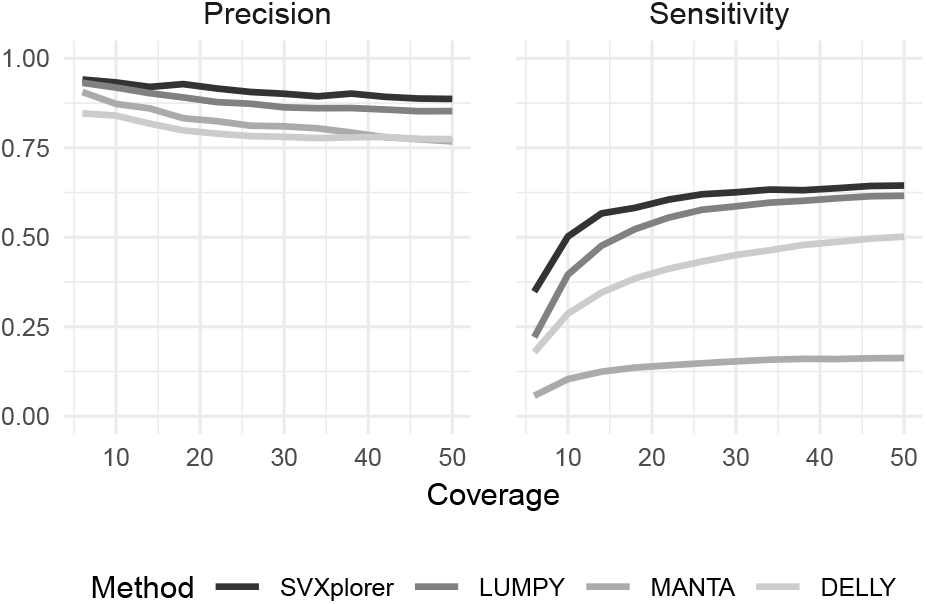
Sensitivity and precision vs coverage for ERR194147

Fig. 4A shows the overall normalized self consistency for the four callers, with SVXplorer showing an improvement of ≈ 5% over the second best. We show plots for each of the three common SV types along with calls by category for each caller in the Supplementary Results, where SVXplorer is the most consistent overall. As alluded to before, the average number of inversions called for the two libraries was 50 for SVXplorer, 30 for LUMPY, 350 for MANTA and 599 for DELLY. SVXplorer and LUMPY are much more in line with expectation(11) compared to DELLY and MANTA.

**Fig. 4.**
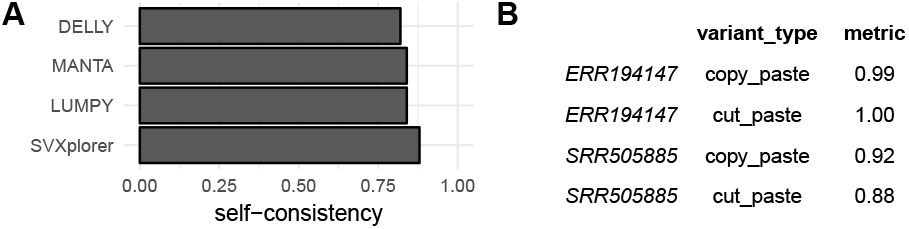
A. Normalized self-consistency comparison for the various approaches. B. Complex variant self-consistency for SVXplorer.

We also evaluated the complex-variant self-consistency for 3-breakpoint complex variants (cut- and copy-paste insertions) for NA12878 using SVXplorer and we report it in Fig. 4B. The 2-breakpoint source location of all insertion calls made by SVXplorer (cut-paste and copy-paste) was extracted for one library and checked for overlap with any “FR”,”RF” cluster or complex variant source location of the unfiltered call set of the other library (see Supplementary Results for details). This check corroborates the correctness of the complex variant breakpoints for a given library via evidence of similar breakpoints in the other library assessed by a simple overlap. The overlap rate being very close to 100% in most cases substantiates that the variants are not products of artifacts in data but real SVs.

#### H.3. AJ Trio

We next evaluated the performance of SVXplorer on the data from the AJ trio sequenced as part of the Genome in a Bottle (GIAB) effort. In general, trio analysis is also useful in testing result reproducibility and accuracy, i.e., we expect that all variants in the child should also be found in the parents and that there must be more variants shared between the child and one of the parents as compared to those shared between both parents. Self-consistency was evaluated as above for NA12878. AJ trio self-consistency for the various callers is shown in Table 3. SVXplorer outperforms the other callers in every category in this analysis – in terms of difference between calls shared between parents and those between child and either parent, in terms of calls found in child but not in either parent, and in terms of raw overlap of calls between child and parent.

**Table 3.**
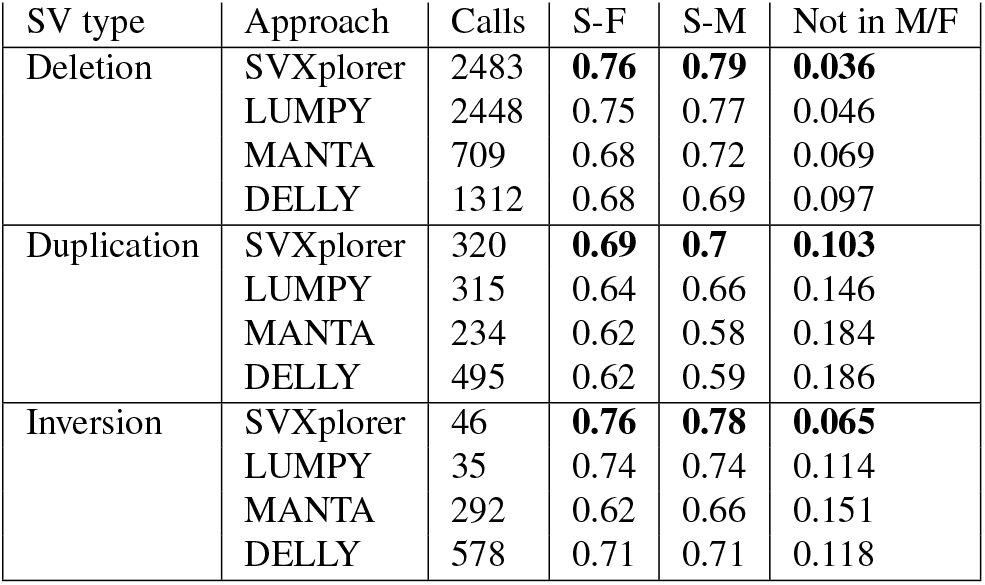
AJ Trio self-consistency: “S-F” refers to the overlap between calls of son and father for each SV category as a fraction of son’s total calls (first column), “S-M” to the same between son and mother, and the last column shows the fraction of the son’s calls that were not seen in either parent. The best result for each metric is highlighted in bold.

We also evaluated the deletion calls for HG002 against an available truth set generated using an ensemble approach in Parliament (9), and show it in Table 4. SVXplorer consistently outperforms the other callers in sensitivity, precision and F1 score. This superior performance further lends credence to various aspects of the self-consistency comparison above.

**Table 4.** Performance of deletion calls for HG002. The best result for each metric is highlighted in bold.

|  | SVXplorer | LUMPY | DELLY | MANTA |
| --- | --- | --- | --- | --- |
| Calls | 2478 | 2448 | 1312 | 709 |
| True-positives | **2057** | 1969 | 795 | 498 |
| Sensitivity | **61.2** | 58.6 | 23.7 | 14.9 |
| Precision | **83.2** | 80.4 | 60.6 | 70.2 |
| F1-score | **70.6** | 67.8 | 34.0 | 24.5 |

## Conclusion

We have developed a structural variant caller that shows improvement over existing approaches on simulated variants and real datasets (haploid and diploid samples). It produces more consistent calls for related individuals as well as for different libraries for the same individual, compared to several other callers. It outperforms compared callers in precision as well as sensitivity, particularly when the coverage is lower or the insert length distribution sharply deviates from a Poisson curve. Unlike most other SV callers, SVXplorer registers deletions and duplications arising from complex variants like translocations and copy-paste insertions, improving the precision of CNVs in the process.

There are several reasons for SVXplorer’s overall effectiveness and better performance. The most significant of those is the pairing of clusters with specified signatures to form above-mentioned 3-breakpoint complex variants, which does not call individual clusters as variants until it has exhaustively analyzed other possibilities. Most callers that rely on paired-end signatures annotate RF clusters as evidence for a duplication and FR clusters as evidence of deletion. Even if read-depth filters are used, the accuracy of such calls can be low, for example in the case of breakpoints generated by retrotransposons that can ‘copy and paste’ their genetic code around the genome. The signature of such calls from discordant reads is an overlapping RF and FR cluster. Without cluster consolidation, a method is likely to call a deletion and duplication in the region with incorrect breakpoints (Fig. 1). SVXplorer’s comprehensive consolidation for insertions arising through ‘cut and paste’ and ‘copy and paste’ mechanisms, inversions, and even tandem duplications enhance its putative call set by reducing false positives among deletions and tandem duplications while identifying accurate, complete insertion sites. Both PE and SR alignments are used individually and collectively to exhaustively form all listed complex variants with specific signatures. The final support thresholds and all other processing are thus applied not to individual clusters but to consolidated variant blocks.

Several enhancements to SVXplorer can be envisioned that would improve its utility and performance. Subsequent to cluster formation, SVXplorer forms breakpoint margins using its insert-length-percentile-based approach, which is not probabilistic. These margins are more precisely handled by LUMPY which uses a probabilistic representation of the SV breakpoint based on the insert length distribution. SVXplorer also does not have an explicit mechanism to identify insertions and deletions smaller than both the insert-length standard deviation and the lowest primary alignment length for the SV. Another area of improvement for SVXplorer is in the handling of multi-allelic variants. For example, a deletion and a duplication with similar reference breakpoints may not be called by SVXplorer as it could be annotated as a copy-number invariant region in the final filter. Such variants, however, can be identified in a family trio by post-processing the identified variants. The current version of SVXplorer does not model biases in sequencing, relying on a careful examination of read-depth instead. However future versions should be able to incorporate better models of read-depth using single-position models, speeding up the execution of the approach.

However, in a more general sense, cluster consolidation which effectively models the smallest set of variants that can coherently be described by the observed signals of PE, SR and read depth provides much improved precision (and sensitivity) in identification of genomic breakpoints. SVXplorer implements that approach for primary alignments and we show an improvement in the precision of the identified variants when compared to several existing callers.

## ACKNOWLEDGEMENTS

This work was supported by NIH award [P30CA044579] to the UVA Cancer Center.

## Supplementary Note 1: Supplementary Methods

### A. Preprocessing

In the sample BAM, only fragments (a) that are marked as concordant by the aligner, (b) where both reads from the fragment align uniquely, and (c) the reads passing a preset alignment-score threshold are used to calculate insert length and coverage.

We filter the BAM file to keep discordant reads that pass preset insert length thresholds relative to the mean (both on *sigma*_3_ as defined in Methods for both positive and negative devations from the mean) and respective mapping quality thresholds. This preset mapping quality threshold is 1 for “FR” clusters and 20 for “RF” clusters, as the latter indicate duplications – which are fraught with surrounding repeats.

### B. Cluster Formation

As part of the cluster formation stage, fragments that seem highly aberrant based on relative left and right tip positions are removed from the cluster in question using a k-means approach. Alignments once finally written in a cluster are not used or counted in subsequent cliques, and clusters with size < *3* are not written by default.

It is worth mentioning here that a “cluster preservation” routine exists for paired-end clusters and can be activated by the user. The routine retains clusters with size < *3* fragments at this point for subsequent stages. If split reads exist in the breakpoint regions of these clusters such that the combined (PE+SR) fragment support for the cluster is greater than the minimum cluster size of 3, the cluster is preserved. This increases run-time for high-coverage data sets but could improve results with low-coverage data.

Several structural variant callers identify regions of the genome where two or more clusters together imply conflicting calls. These conflicts are caused by 3 variants existing successively within each other, such as 2 deletions and a non-deletion, 2 inversions and a non-inversion etc. SVXplorer did not detect any conflicts of this nature. We impose a more stringent criterion and identify regions that contain multiple clusters with the same orientation close to each other. Such regions are typically indicative of misalignments. We store the coordinates for such clusters in a separate file, and these regions are subsequently processed to ascertain if any of the clusters can be selected based on a predefined threshold of proportion of alignments in the region it accounts for, and included in subsequent analyses.

### C. Cluster Consolidation

As referenced in the manuscript, copy-paste and cut-paste insertions are shown and described in more detail in Figures S1 and S2.

**Fig. S1.**
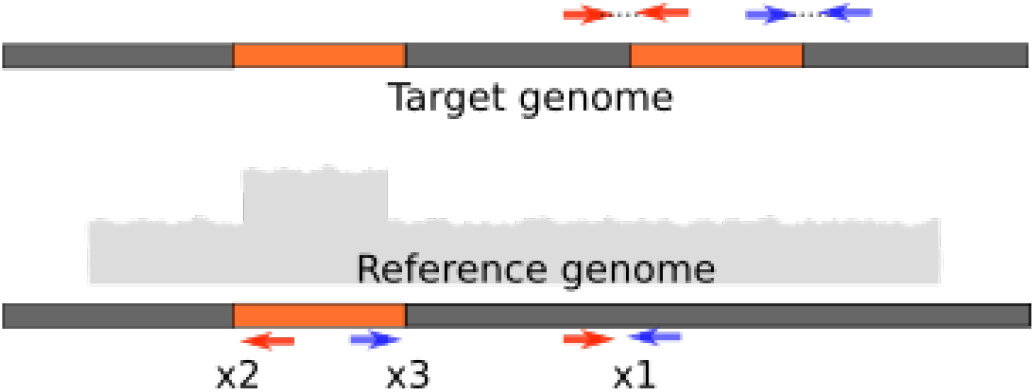
A simple copy-paste insertion. The segment in orange is duplicated downstream in the sample. The figure shows 2 distinct clusters in red and blue matching up in the reference to form a copy-paste insertion. Breakpoint 1 (*x*_1_) is defined to be the overlap of adjacent oppositely-oriented alignments from the 2 clusters, and breakpoints 2 and 3 (*x*_2_ and *x*_3_) are defined by their respective mate alignments, with *x*_2_ < *x*_3_ by convention, whether upstream or downstream from *x*_1_.

**Fig. S2.**
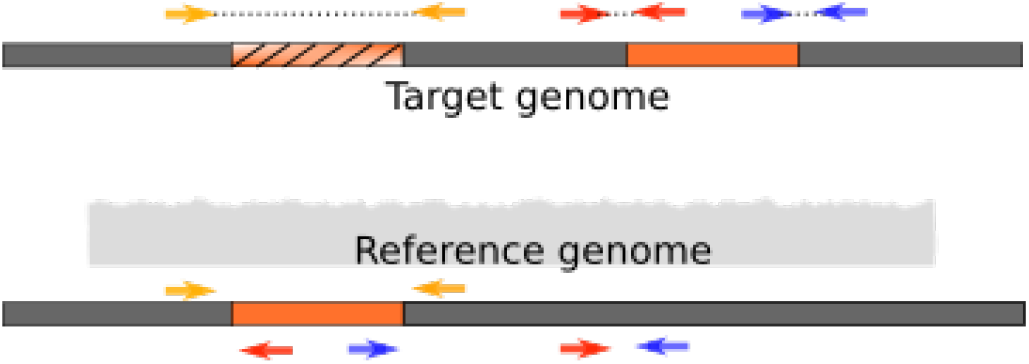
A simple cut-paste insertion (translocation). The segment in orange is deleted and pasted downstream in the sample. The figure shows 3 distinct clusters, shown in red, blue and light orange. The cluster shown in light orange is the extra “FR” cluster resulting from the deletion of the translocated segment

The VCF file groups multiple events (DUP, BND) coming from one cut-paste or copy-paste insertion via the GROUPID subfield (in “INFO”) in order to preserve all the information of the BEDPE output. It also contains a “comment” subfield where the likely but undetermined SV type of a BND event is printed. An “ISINV” flag is printed in the “INFO” field if a duplication is inverted.

SVXplorer attempts to account for many special cases of SV formation. One worth mentioning here is the “crossover” TD cluster. As shown in Fig. S3, such a cluster is formed when the insert length is comparable to the size of the tandem-duplicated segment. The cluster consists of paired-end alignments whose reads are aligned as “FR” (but are very close or may even overlap each other). Therefore, the left breakpoint of the cluster is defined by reverse-stranded alignments and the right breakpoint is defined by forward-stranded alignments. This is unlike a deletion “FR” cluster and is processed as a tandem duplication. It is also of note that a small “FR” cluster can be formed around any of the above variant types that involve an inserted segment (i.e., tandem duplication and all insertions) due to the insert length of the aligned segments being less in the reference than in the sample. Thus, such a cluster can be part of any of the above signatures involving inserted segments.

**Fig. S3.**
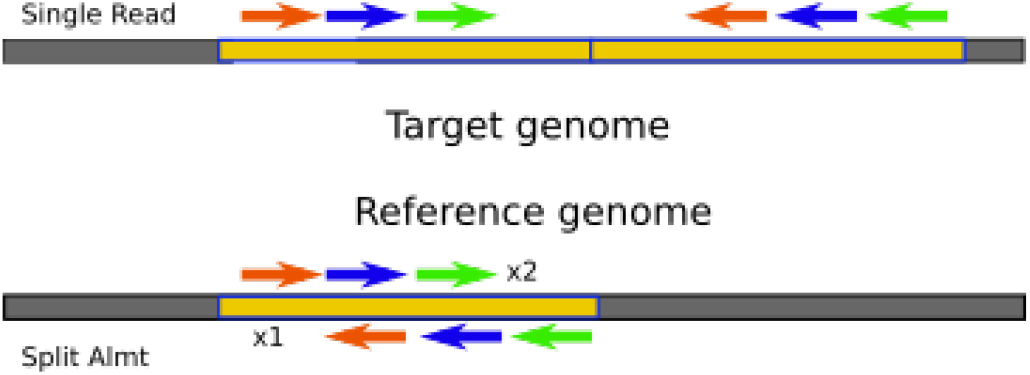
A “crossover” TD cluster. The segment in yellow is adjacently duplicated downstream in the sample. The figure thus shows sequenced fragments from a tandem duplication that align as “FR.” In such a case, the left breakpoint is defined by reverse alignments and the right breakpoint is defined by forward alignments.

### D. Incorporation of split-reads

If the alignment signature supports an existing variant and the split alignments have breakpoints within existing breakpoint margins (see Fig. S5), then we use the read alignment in the reference to update the current variant breakpoints and tag the variant as “precise”. The alignment record is added to the existing variant set in the variant map, and the variant is tagged as “PE_SR”-supported. For example, an “FF” PE cluster may now be supported by an “FR” SR cluster, and if it happens to join the reference on the other side of the potential inversion as the PE cluster, then it completes the putative inversion (there is a “liberal inversion” parameter that can be set by the user to merely require support for inversions by PE and SR reads to be called). In comparing alignments with existing SR variants a small bidirectional “slop” is used to account for possible imprecision in some reads (5 by default).

Variants called exclusively by split-reads are more limited in their scope compared to PE variants for Illumina reads. SR coverage is typically lower than PE coverage, which may render SR cluster matching and consolidation less reliable considering the narrower Poisson window for such reads in sequencing (reads overlapping variant boundaries, as opposed to the more numerous whole reads outside boundaries supporting the variant). For example, a single split read may be aligned such that its two split partners align with the same orientation (“FF” or “RR”) in the reference. This could indicate a deletion or “two-thirds” of a copy-paste insertion or translocation (see Fig. S4). Thus, if an “FF” or “RR” split-read cluster is unmatched till the end, it is not labelled a putative deletion immediately but only if it passes stringent deletion filters in the final pile-up filtering stage as described below.

**Fig. S4.**
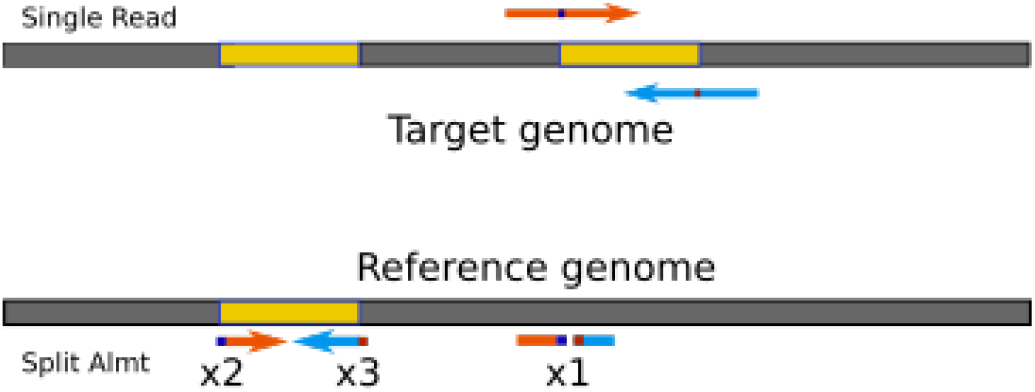
A copy-paste insertion call from split reads. The segment in yellow is duplicated downstream in the sample. The orange read by itself would lead to a TD_I call and the blue by itself to a DEL_INS call. But together they define a copy-paste insertion consisting of 3 distinct breakpoints.

If the split partners are swapped, then they could indicate a tandem duplication or a copy-paste insertion/translocation. A *swapped* alignment is defined as a split read where one split partner sequence came before the other in the sequencing direction in the sample, but now comes after the other in the alignment direction in the reference (as in Fig. S6). The swap is determined by extracting the relative query start and end positions from the BAM file (applicable for all “FR” and “RF” alignments). New SR variant categories are: deletion/insertion, tandem duplication/insertion, insertion and inversion. A brief description of these signatures is provided now.

**Fig. S5.**
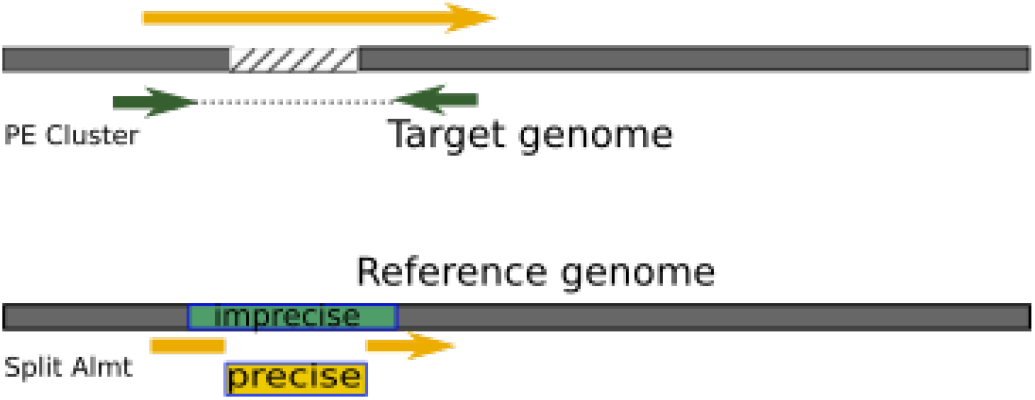
Example of a PE deletion call supported by a split read. The read shown in yellow (size exaggerated in target) is split into 2 alignments in the reference close to the PE breakpoints. The segment in green is the putative PE deletion call and the segment in yellow shows revised precise breakpoints.

**Fig. S6.**
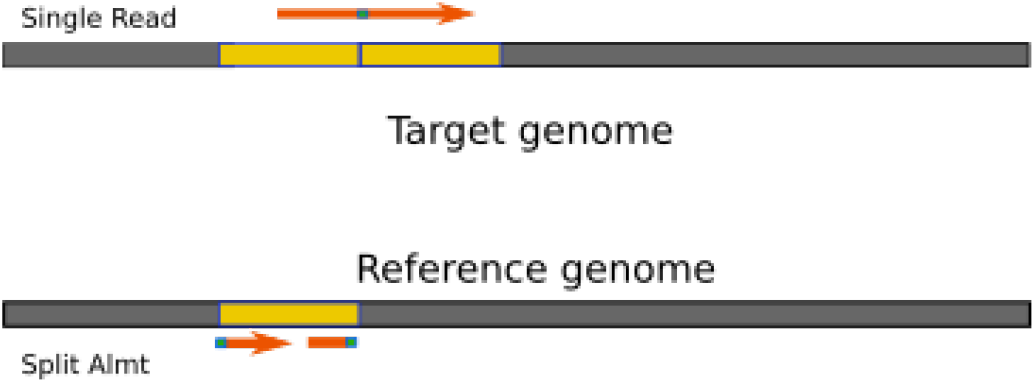
A TD_I call from split reads. The segment in orange is tandem-duplicated downstream in the sample. The read shown in orange splits in alignment at the point shown in blue. The split partners are swapped in alignment, i.e., the head portion of the original forward-oriented read aligns in the reference to the left of the tail portion of that read. Such cases give rise to a TD_I cluster.

The requirement for both ends of an SR-supported inversion (as with PE and mixed inversions) to join the reference is to substantiate the existence of an actual inversion, as there may be other kinds of inverted structural variations or artifacts present.

### E. Variant Filtering

For our case of primary alignments, variant filtering particularly addresses complex variants that have one cluster in common and the other cluster unique to each variant (see Fig. S7). This is indicative of an atypical situation with regard to variant identification and therefore neither variant would be called unless one of them exceeds the support threshold. If both exceed the support threshold, they are addressed in the next stage. Such situations can be often seen in real sequenced samples, but are revealed upon visual analysis as artifacts. Variants that are not supported by expected clusters, for example, an inversion that is only supported by a “FF” cluster are not trustworthy as calls in any particular variant category and included simply as BND events. Also, a cluster that may have been too small to make the final support threshold filter by itself but fits cogently into an existing variant with sufficient support would now be counted and useful.

To calculate the disjointness threshold, RSVSim was used to alter the “hg19” human reference genome by introducing 500 deletions, tandem duplications, inversions and insertions (i.e., translocations and copy-paste insertions) and we used wgsim to synthetically read from this sample at coverages from 5x to 45x in steps of 10, and with a standard deviation ranging from 10 to 70 in steps of 20. SVXplorer was run on these different data sets with different minimum thresholds for support (in three different variant categories: PE, SR and “mixed” or PE+SR) and the F1 score for the identified variants was calculated against the true variant set. The threshold yielding the highest F1 score was recorded for each data set as a function of (coverage, insert length std) and used to generate a best-fit line, and all intermediate coverage values are fit by linear interpolation. The dependence on standard deviation in insert length ends up being inconsequential in this simple model.

**Fig. S7.**
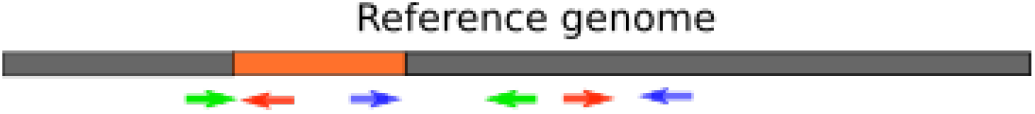
A case where 2 PE clusters each separately match up with a third cluster. The clusters in red and green match up with each other and so do the ones in red and blue, each matching pair indicating a copy-paste insertion. It is quite unlikely that both are true. This is addressed in the filtering stage.

### F. Incorporation of depth of coverage

Coverage for each chromosome or contig is calculated using mappable regions only. All final SV calls are also made using coverage calculated in the variant region using mappable bases only. Coverage calculation seeks to use other bases only if a predefined sufficient number of mappable bases is not seen. Only uniquely aligning reads are used in calculation of coverage and reads that could be putative PCR duplicates, or refer to secondary alignments, are filtered away.

Variant-region coverage information is recorded in the INFO filed in the VCF file if a certain variant was rejected by unexpected read depth. This filter is used to enhance calls made in all SV categories except inversions.

Variants that are supported exclusively by SRs and called as possible deletions, tandem duplications or copy-paste insertions are rejected if contradicted by preset thresholds. Deletion and duplication calls that are well supported by PE alignments are not required to satisfy the preset thresholds but are not written if the VCR exceeds or falls below a slightly more liberal rejection threshold. This is because, as alluded to in an earlier section, an unmatched “FR” PE cluster that is above the coverage-determined support threshold and composed of fragments with large insert lengths is more likely a deletion than an “FF” or “RR” SR cluster.

Additionally, some routines are used to analyze the coverage of complex variants (such as cut-paste and copy-paste insertions) in regions between relevant pairs of breakpoints and decouple them into BND events if necessary, if the SPLIT_INS parameter is turned on. As an example one “FR” and and one “RF” PE cluster may have combined to form a copy-paste insertion, but if it is now seen that the coverage between the “FR” breakpoints indicates a deletion, then the insertion call is broken into two simple BND events to be safe.

## Supplementary Note 2: Supplementary Results

### A. Simulated Data

As expected, none of the tools made substantial false-positive calls at coverages higher than 6X (SVXplorer was uniformly the leader by a small margin ranging from .1 – .3%). This is shown in Fig. S8.

**Fig. S8.**
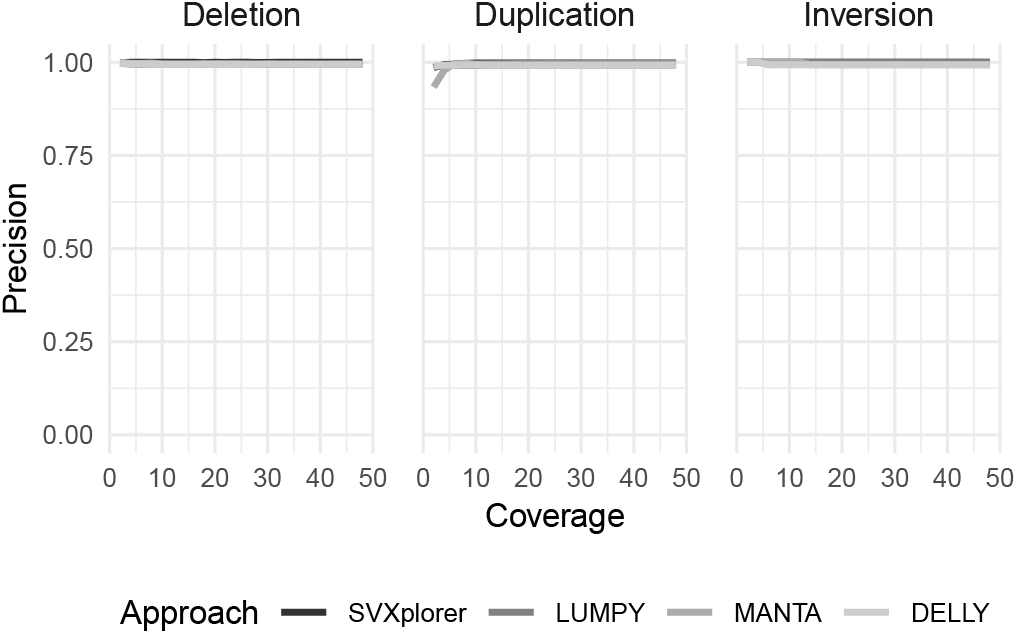
Precision vs coverage for simulated data

### B. CHM1

We reevaluated the various performance metrics for the deletion calls made for CHM1, this time stratified based on the genome annotations in those regions. Fig. S9 shows a detailed relative analysis by category. SVXplorer has the highest sensitivity for 14 of the 17 different region categories, whereas LUMPY had the highest detected sensitivity for 7 of the 17 annotated categories. Statistics are too sparse (in the number of true calls and number of catches in general) in CHM1 inversions to have a meaningful relative comparison of the approaches. Insertion locations were not assessed as most tools neither call de novo insertions nor include explicit paste locations for other insertions.

### C. NA12878 Self-Consistency

As seen in Fig. S10, all the callers seem to perform reasonably well on self-consistency. SVXplorer is generally best or the second-best in each category and obtains the highest normalized (by number of each SV type) self-consistency (by 5% over the second-best) with 6867 calls (LUMPY has 7012, DELLY has 6708 and MANTA has a total of 2827 SV calls). The average number of inversions called for the two libraries was 50 for SVXplorer, 30 for LUMPY, 350 for MANTA and 599 for DELLY. SVXplorer and LUMPY are much more in line with expectations(12) compared to DELLY and MANTA.

In the AJ-Trio analysis, similar to the idea in NA12878 with sequencing libraries, we are interested in knowing whether a variant found in the child presents *any* evidence of being seen in either parent

### D. NA12878 Complex Variant Self-Consistency

The purpose of this simple check was to see if the complex variants source breakpoints (which are usually defined by multiple clusters) are seen to at all overlap either with an “FR” or “RF” cluster, or another complex variant source location, in the other library. This method to assess complex overlap was chosen because very often only partial signatures of complex variants occur (e.g., only one “FR” or one “RF” cluster for a cut-paste or copy-paste insertion) due to reasons related to coverage, homology, repeats and alignment. Therefore, if most source locations of one library overlap with another similar location in the other library, it is a good indicator of self-consistency for complex variants. As shown in Fig. 4, SVXplorer has 100% (8 of 8) and 89% (8 of 9) self-consistency for cut-paste insertions for the two libraries whereas for copy-paste insertions it has 99.5% (191 of 192) and 91% (133 of 146). The difference in self-consistency for the two libraries arises due to the fact that the sequencing, insert length distribution etc. for the two are very different. The “SRR505885” library has many more lower mapping-quality calls, particularly of “RF” alignments.

### E. AJ Trio

We see that the percentage of calls that were in the son and not found in either parent was lowest for SVXplorer for each variant type: 3%, 10% and 7% for deletions, duplications and inversions, respectively, whereas the second best numbers (not necessarily by the same caller) are 5%, 15% and 11% respectively. All callers perform as expected when evaluating the difference between calls shared by the child and one of the parents and those shared by both parents, with the difference never falling below 10% of the child’s calls for any variant type. The number of inversions called by SVXplorer (average = 44, on same order as LUMPY’s 30) is supported by other reports (11) whereas MANTA and DELLY call 288 and 556 respectively. The lower number of inversions is effected by SVXplorer’s requirement to see both ends of the inversion joining the reference, by a combination of PE or SR reads. For reference, SVXplorer makes a total of 2802 calls for the son, LUMPY makes 2798, DELLY makes 2385 and MANTA makes 1235 calls. AJ trio self-consistency for the various callers is shown in Fig. S11.

**Fig. S9.**
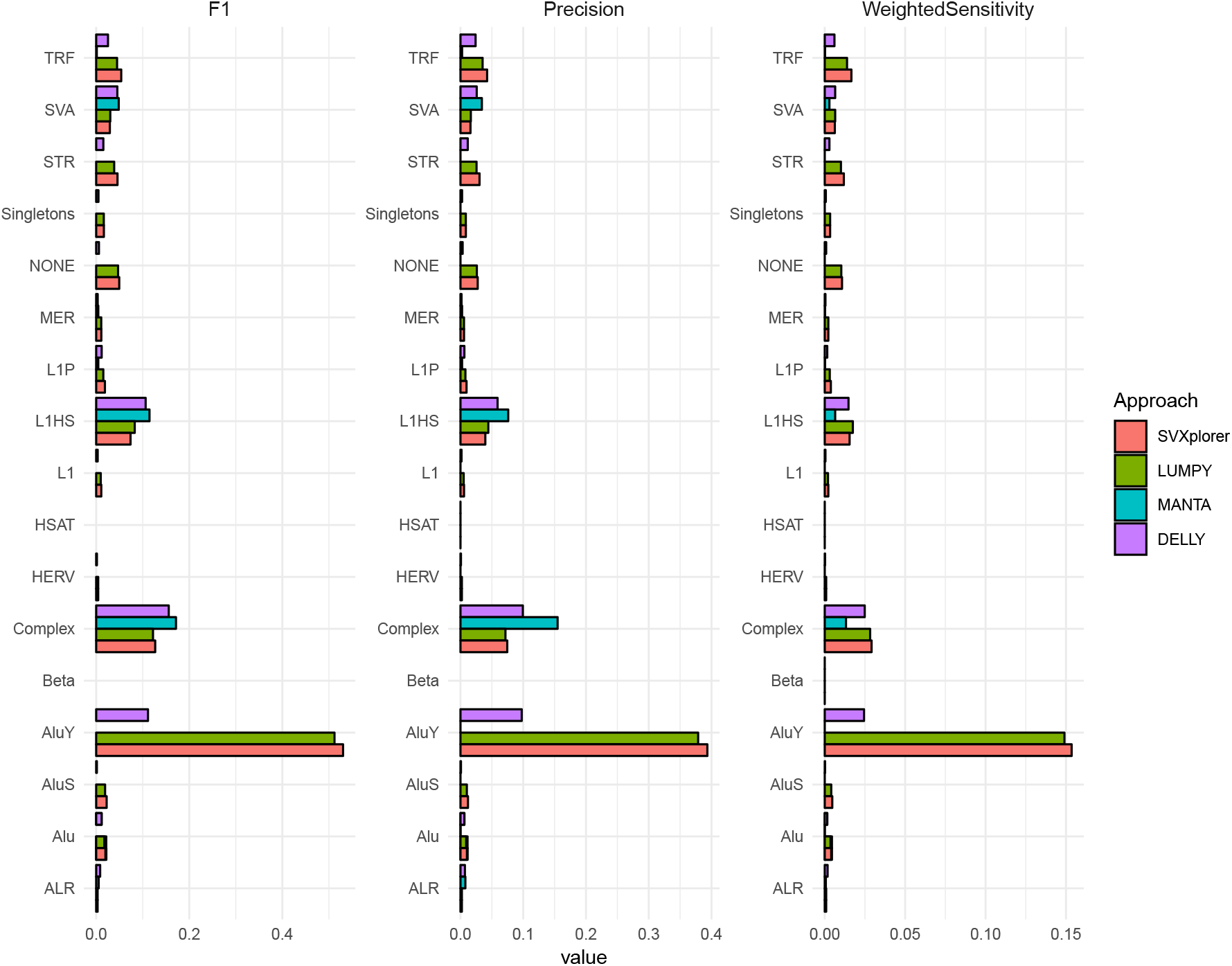
Evaluation metrics for CHM1 Deletions using genome annotations

**Fig. S10.**
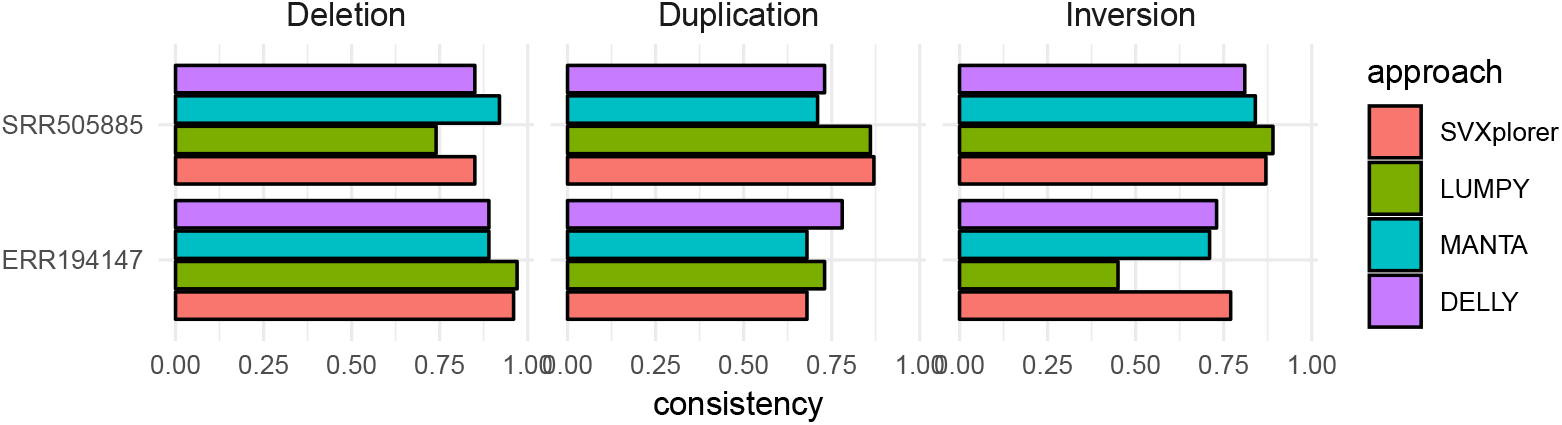
Self-consistency in NA12878 data when various approaches are used. “Consistency” refers to the fraction of calls in the listed base library that were found in the other library.

**Fig. S11.**
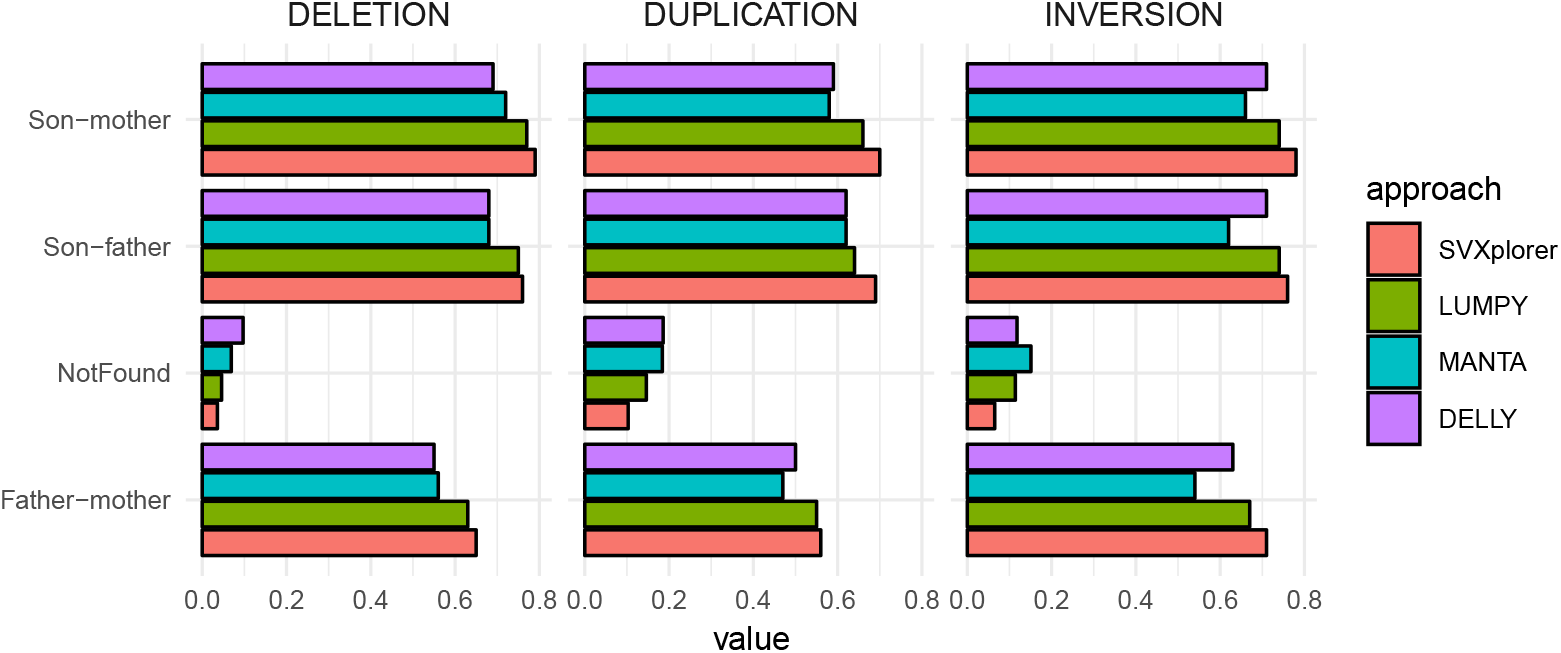
AJ Trio self-consistency for the various SV types. “A-B” refers to fraction of total calls made for A that were found in B. Here A or B is a placeholder for either son, father or mother. “Difference” refers to the difference between calls in common between both parents and those in common between son and parents (normalized). We expect this to be large. The “not found” column shows the fraction of total calls that were made in the son that were not found in either parent.

## Footnotes

1 The calculated insert length accounts for read orientation in reference

2 The denominator is also somewhat dependent but, given the spread/smattering of discordant alignments in the genome, it has opposite monotonicity to the numerator and only supports the same monotonic behavior. Thus, it need not be further treated or considered for this heuristic motivation of the connection weight

3 A typical SVXplorer workflow takes ≈ 17 minutes on a 30X simulation

